# Crossing thermal limits: functional collapse of the surfgrass *Phyllospadix scouleri* under extreme marine heatwaves

**DOI:** 10.1101/2025.11.12.688107

**Authors:** Manuel Vivanco-Bercovich, Paula Bonet-Meliá, Nadine Schubert, Lázaro Marín-Guirao, Raquel Muñiz-Salazar, Alejandro Cabello-Pasini, Alejandra Ferreira-Arrieta, Jessica Anayansi Garcia-Pantoja, José Manuel Guzmán-Calderón, Gabriele Procaccini, Guillermo Samperio-Ramos, Jose M. Sandoval-Gil

## Abstract

Marine heatwaves (MHWs) are intensifying under climate change, yet the physiological limits that constrain seagrass resilience remain poorly defined. We experimentally tested the responses of the surfgrass *Phyllospadix scouleri*, a foundation species of the Northeast Pacific coast, to simulated MHWs of contrasting intensity. In a 27-day mesocosm experiment, plants were exposed to fluctuating temperatures representing a severe MHW (23.5 ± 1.5 °C) and an extreme MHW (26.5 ± 1.5 °C), while photosynthetic performance, respiration, nitrogen metabolism, oxidative stress, and growth were monitored during and after warming. *Phyllospadix scouleri* maintained photosynthetic capacity and carbon balance under severe warming but exhibited pronounced physiological disruption at extreme temperatures, including sustained photoinhibition, reduced nitrate assimilation, elevated respiration, and negative daily productivity. These effects persisted after heat stress, leading to reduced growth and indicating incomplete recovery. Multivariate analyses revealed a distinct transition from tolerance to functional breakdown near 26.5 °C, suggesting a physiological tipping point only 5–6 °C above current summer maxima in the area of the studied population. Our findings demonstrate that intensifying MHWs may rapidly erode the thermal safety margin of temperate seagrasses, pushing foundational coastal ecosystems toward metabolic instability and potential regime shifts under continued ocean warming.

**Highlight:** Extreme marine heatwave disrupts photosynthesis, nitrogen metabolism, and carbon balance in the seagrass *Phyllospadix scouleri*, suggesting a narrow thermal safety margin in the face of ocean warming.

## 1. Introduction

Seagrass meadows are declining globally at alarming rates due to growing pressures from climate change and human activities (Dunic *et al*., 2021). As foundation species, their degradation implies the loss of entire ecosystems, with far-reaching ecological, socio-economic, and cultural consequences that threaten fisheries, shoreline stability, blue-carbon storage, and multiple societal values (Unsworth *et al*., 2022, Foster *et al*., 2025). Among the most severe disturbances affecting seagrass meadows is ocean warming, which is expected to drive a global spatial redistribution of seagrass species (Gouvêa *et al*., 2024). Closely associated with this long-term trend are marine heatwaves (MHWs), defined as discrete periods of anomalously high sea surface temperature relative to the local climatology (Hobday *et al*., 2016), which can trigger large-scale degradation and even mass mortality events in seagrass meadows (e.g., Marbà & Duarte 2010, Strydom *et al*., 2020).

Current knowledge on seagrass thermal physiology spans many species worldwide, yet remains dominated by studies on temperate taxa such as *Posidonia oceanica*, *Zostera marina*, and *Cymodocea nodosa* (Table S1). These experiments have tested warming scenarios ranging from +2 to +20 °C above local climatology and lasting from a few hours to several months. Although many exceed realistic MHW conditions, they consistently show that elevated temperatures reduce photosynthetic efficiency, deplete photosynthetic pigments, and increase oxidative stress, leading to reductions in growth, biomass, and shoot density. When thermal thresholds are surpassed, it results in irreversible tissue damage, necrosis, and mortality. Altogether, this evidence demonstrates the nonlinear thermal sensitivity of seagrasses, with sharp declines beyond species-specific limits.

These temperature-induced impacts arise from the disruption of fundamental physiological processes, including photosynthesis, oxidative balance, and carbon allocation (Nguyen *et al*., 2021). The tolerance of seagrasses to elevated temperatures is modulated by intrinsic biological attributes at both species and population levels, such as acclimative plasticity and adaptive capacity (Pazzaglia *et al*., 2021). Moreover, thermal tolerance can be further influenced by factors such as local adaptation, seasonal timing, plant ontogeny, and tissue age (Ruocco *et al*., 2019; Beca-Carretero *et al*., 2021; Entrambasaguas *et al*., 2021; Rinaldi *et al*., 2023). Evaluating thermal sensitivity across populations and temperature gradients is therefore essential to refine forecasts of seagrass vulnerability under future climate scenarios (Bennett *et al*., 2019; Starko *et al*., 2024).

Seagrass responses to warming also depend largely on the characteristics of each MHW, particularly its intensity, duration, and frequency. Moderate temperature increases may transiently stimulate metabolic processes such as photosynthesis (Beca-Carretero *et al*., 2018), whereas more intense or prolonged events exceed species’ thermal limits, causing oxidative stress, reduced growth, and mortality (Collier & Waycott, 2014). As climate change progresses, MHWs are becoming more frequent, longer lasting, and more intense, with “extreme” events - once rare - projected to represent up to 70% of all occurrences by 2100 (Oliver *et al*., 2019). To better anticipate their ecological consequences, it is increasingly important to assess seagrass responses across warming levels and under temperature regimes that mimic, as closely as possible, real MHW conditions, encompassing both contemporary events and projected future scenarios (e.g., Saha *et al*., 2020; Stipcich *et al*., 2022a; Gillis *et al*., 2025).

Along the Northeast Pacific, and particularly on the coasts of the United States and Baja California (Mexico), the frequency and intensity of MHWs have increased markedly in recent decades (Bond *et al*., 2015; Wei *et al*., 2021). These extreme events have triggered widespread ecological disruptions, including persistent declines in macrophyte assemblages such as giant kelp forests (*Macrocystis pyrifera*; Arafeh-Dalmau *et al*., 2019; Beas-Luna *et al*., 2020) and associated shifts in community structure. While much attention has been focused on canopy-forming kelps, other neighboring coastal primary producers may be equally vulnerable to increasing ocean temperatures.

Surfgrasses (*Phyllospadix scouleri* W.J. Hooker and *Phyllospadix torreyi* S. Watson) are key foundation species that dominate the intertidal and shallow-subtidal rocky shores of the Northeast Pacific, from Vancouver Island (Canada) to Baja California (Mexico) (den Hartog & Kuo, 2006). Their high productivity (> 8000 g DW m⁻² yr⁻¹; Ramírez-García *et al*., 1998) and dense canopies (8000–12000 shoots m⁻²; García-Pantoja *et al*., 2020) create structurally complex habitats that support diverse assemblages of invertebrates and fish. With leaves that can exceed 2 m in length, surfgrass meadows modulate physical, chemical, and biological processes across intertidal pools and shallow (∼10 m) rocky bottoms (Tharaldson, 2018; Ruiz-Montoya *et al*., 2021). *Phyllospadix* meadows provide essential ecosystem services, including coastal protection, nutrient cycling, carbon sequestration, and habitat provision (García-Pantoja *et al*., 2020; Fields & Silbiger, 2022). Unlike most seagrasses, however, surfgrasses thrive on wave-exposed rocky substrates, an adaptation made possible by their specialized rhizome anchoring system (Cooper & McRoy, 1988), which enables them to colonize high-energy coastal zones often inaccessible to other vascular plants.

Despite their ecological importance, surfgrasses remain among the least studied seagrass groups worldwide (Strydom *et al*., 2023). Early work on *P. torreyi* identified an optimal growth range of 12–14 °C and sharp declines above 17 °C, indicating a relatively narrow thermal tolerance (Drysdale & Barbour, 1975). At lower latitudes, recent experiments revealed that *P. torreyi* can transiently increase photosynthetic capacity under short-term heat exposure (∼25 °C for seven days), although this response coincides with carbon reserve depletion, suggesting energetic trade-offs (Vivanco-Bercovich *et al*., 2022). In *P. scouleri*, repeated exposures to warming (∼24 °C) led to progressive declines in photosynthetic efficiency, oxidative stress accumulation, and reduced nitrate uptake, despite temporary maintenance of growth through carbon mobilization (Bonet-Melià *et al*., 2023; Vivanco-Bercovich *et al*., 2024). Intertidal populations appear relatively tolerant to elevated temperatures (Menge *et al*., 2020; Ruiz-Montoya *et al*., 2021), but recent findings indicate that climate change is already shaping population structure and genetic diversity across *Phyllospadix* species (Tavares *et al*., 2024). Collectively, these studies show that surfgrasses can initially withstand moderate warming, but major uncertainties remain about the physiological thresholds and resilience of *P. scouleri* under the extreme warming scenarios projected for coming decades.

This study aimed to compare the physiological responses and growth of *P. scouleri* under MHWs of different intensities, representing contemporary and future scenarios. We conducted a 27-day mesocosm experiment in which subtidal adult plants were exposed to two scenarios: (i) a severe MHW, with a maximum intensity of +5 °C, representing contemporary events, and (ii) an extreme MHW, reaching +8 °C, simulating future projections. Daily temperature fluctuations were incorporated in the experiment to resemble natural conditions. Across two experimental phases (during heat exposure, and after warming cessation), we analyzed a set of ecophysiological descriptors to identify acclimation mechanisms and physiological responses. These findings provide insights into the potential thermal thresholds of *P. scouleri*, offering a better understanding of its capacity to withstand climate-driven heat extremes, and supporting conservation strategies for these ecologically and functionally important coastal ecosystems.

## 2. Material and methods

### 2.1. Donor meadow and thermal history

The donor meadow is located on Todos Santos Island (31° 48’ 25.66” N, 116° 47’ 46.41” W), approximately 19 km offshore from Ensenada, Baja California, México, within the Pacific Islands Biosphere Reserve (Fig. 1A, B). This meadow spans over >32.000 m^2^, extending from the intertidal to depths of ∼ 7 m along a high energy, rocky coastline (Garcia-Pantoja *et al*., 2020). Seawater temperature in this region typically ranges between 15°C to 20°C, with peaks usually occurring in August - September (Fig. S1A). Yet, in the last few years, summer peak temperatures have repeatedly surpassed 23°C (Fig. S1B).

**Figure 1.**
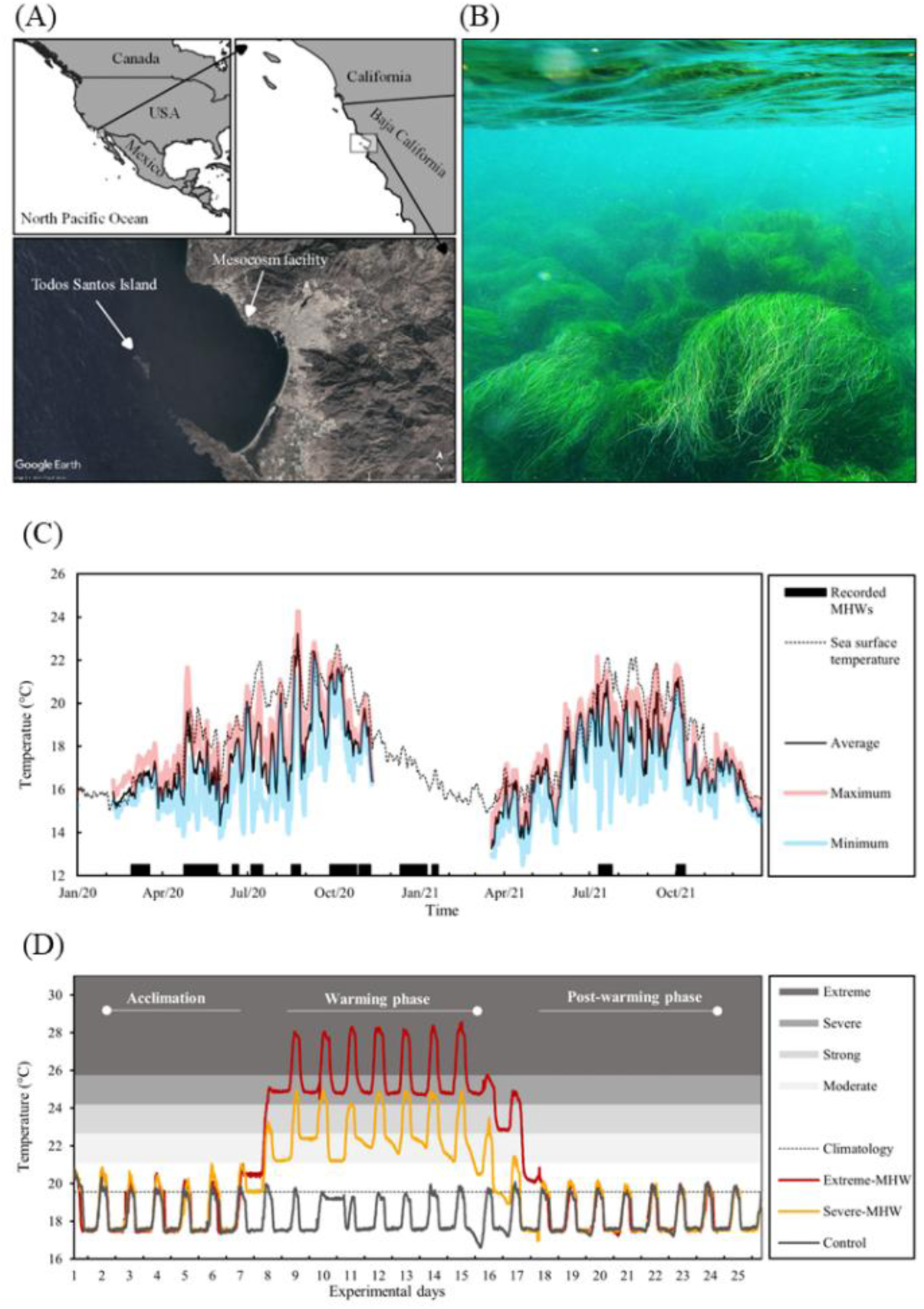
(A) Location of the *Phyllospadix scouleri* donor meadow at Todos Santos Island, Baja California, Mexico, and (B) representative photo of the meadow. (C) Seawater temperatures recorded over two years using submersible loggers at 15-minute intervals. Solid lines show daily average (black), maximum (red), and minimum (blue) temperatures. Satellite SST data (dotted line) were retrieved from NOAA (High-Resolution SST data), and MHWs (black bars) were identified using the Marine Heatwave Tracker (Schlegel et al., 2020). (D) Experimental temperature profiles for Control, Severe-MHW, and Extreme-MHW treatments across the acclimation, warming, and post-warming phases. Sampling time points for biological descriptors are indicated with white dots. Background shading corresponds to MHW categories based on climatological thresholds (Hobday et al., 2018).

Historical satellite-derived sea surface temperature (SST) data indicate that this region has experienced over 100 marine heatwaves (MHWs) since 1982 (Marine Heatwave Tracker platform, www.marineheatwaves.org; Pixel: 116.875°W 31.875°N; Fixed baseline: 1982-2011). Most of these events lasted 10 days or less and were characterized by intensity anomalies ≤ 3°C, although a few reached intensities of up to 5.5 °C (Fig. S2). According to the classification framework proposed by Hobday *et al*. (2018), 88% of the events were categorized as “moderate”, 18% as “strong”, while only three were classified as “severe” and one as “extreme”.

Submersible sensors (Onset-HOBO MX2202) were calibrated and anchored for two years within the donor meadow to obtain high-resolution, sub-daily thermal records beyond what is provided by daily mean satellite SST data. In situ temperature data from sensors showed high thermal variability at the donor meadow (Fig. S3). Discrepancies between satellite-derived and *in situ* measurements exceeded 3°C, with satellite SST underestimating maximum temperatures by ∼2°C. For instance, *in situ* sensors recorded a peak of 24.3°C in August 2020, which was 1.7°C higher than satellite-derived values and nearly 4°C above the climatological mean (Fig. 1C). This dataset was used to define the experimental temperature regimes, ensuring that treatments reproduced both the daily thermal variability and the magnitude of MHWs observed in the field.

### 2.2. Plant collection and acclimation

*Phyllospadix scouleri* plants were collected in September 2021 by divers from the subtidal zone (∼5 m below mean sea level) of the donor meadow. Subtidal plants were selected over intertidal individuals because they are typically exposed to less extreme temperatures and less pronounced daily temperature fluctuations (Ruiz-Montoya *et al*., 2021), and therefore likely represent the portion of the meadow most sensitive to MHWs. To preserve clonal integrity, plants were carefully detached from the rocks while maintaining ramet connections. Samples were transported in coolers within 2 hours to the Instituto de Investigaciones Oceanológicas (IIO) at the Universidad Autónoma de Baja California (UABC). To characterize field physiological conditions prior to acclimation, photobiological descriptors were measured immediately after collection, and leaf tissues were frozen at −80°C for subsequent biological analyses (Table S2).

Plants were acclimated for five days in an outdoor experimental system at a seawater temperature of 18.5 ± 1.5°C, consistent with the thermal regime recorded during the three weeks preceding collection (Fig. 1C). This system consisted of 12 independent tanks (1000 L), each serving as an experimental unit (EU). Each EU contained 6-8 shoot clumps (with ∼200 connected shoots each), which were secured to rocks at the bottom of the tanks. To replicate natural light conditions at the collection site, neutral shading screens were used in order to maintain a maximum light intensity of ∼200 μmol photon m⁻² s⁻¹ at midday, and a daily PAR dose of approximately 4.5 mol photon m⁻² day⁻¹. Photosynthetically active radiation (PAR) and seawater temperature were monitored during the experiment both in the field and in the tanks using submersible data loggers (Onset HOBO MX2202) (Fig. 1D).

### 2.3. Thermal treatments: rationale and experimental setup

The experiment consisted of three temperature treatments: a control (18.5 °C), a severe-MHW (23.5 °C), and an extreme-MHW (26.5 °C) (Fig. 1D). Daily fluctuations of ± 1.5 °C were applied across all treatments to mimic natural diel variability recorded *in situ* (Fig. S3). The control was set to match the temperature recorded at the meadow at the time of sampling, approximately 1 °C below the climatological temperature for that time of year (i.e., 19.5 °C, Fig. 1D).

The severe-MHW treatment (with peaks of 25 °C) represents a realistic contemporary anomalous event based on recent summer observations (Fig. S2). The extreme-MHW treatment was designed to simulate a plausible upper-bound scenario under end-of-century warming. According to the IPCC (2021), sea surface temperatures in the North Pacific are expected to increase by approximately 3 °C by 2100 under high-emission scenarios (SSP5-8.5). Furthermore, MHWs are expected to become more intense as baseline temperatures rise and ocean–atmosphere feedbacks amplify thermal anomalies (Frölicher *et al*., 2018; Oliver *et al*., 2019; Cael *et al*., 2024; Athanase *et al*., 2024). Combined with local sub-daily variability, these projections suggest that short-term temperature extremes in the range of 26.5 ± 1.5 °C are increasingly to occur in upcoming decades. The classification of the simulated MHWs followed the criteria proposed by Hobday *et al*. (2018), which define intensity levels based on peak temperature anomalies relative to local climatology.

Each MHW treatment lasted seven days at peak temperature, flanked by two-day warming and cooling transitions (Fig. 1D). During these transitions, the temperature increased or decreased at rates of ∼2 °C day⁻¹ for the severe-MHW and ∼4 °C day⁻¹ for the extreme-MHW, values within the range of daily variation recorded by in situ sensors at the study site. Following the heatwave period, all plants underwent a 7-day post-warming phase at control temperature (18.5 ± 1.5 °C), identical to that at the control treatment. Physiological responses were assessed at two time points: at the conclusion of the heatwave period (warming phase) and seven days after temperatures returned to control levels (post-warming phase).

Within the mesocosm facility, temperature was regulated using chillers (Aqua Logic Multi-temp Chiller MT-4, 3HP) and submersible heaters. A water pump (AQUAPAK SUPRA 15/1230, flow rate: 100-390 L/min) was utilized to recirculate seawater between reservoir tanks and the four EUs of each treatment. Aeration was supplied from the bottom of each tank to ensure vertical mixing and water column homogenization. Partial seawater replacements were made during the experiment to maintain water quality. Salinity was kept at 33.4 ± 0.3. Distilled water was added daily to offset the loss of water through evaporation.

### 2.4. Physiological traits and growth

At the end of the two experimental phases, random shoots were collected from the EUs for physiological analysis and growth measurements. For each physiological trait, measurements from two samples per EU were averaged to obtain an accurate experimental replicate (n = 4 per treatment), except for photosynthesis-irradiance (P-E) curves, which were conducted on a single sample per EU. The same number of replicates was utilized for the initial condition characterization. To standardize the measurements, all samples were taken from the middle section of the second mature leaf.

#### 2.4.1. Photosynthesis and Respiration (P vs. E curves)

Photosynthetic and respiratory rates were measured in leaf segments using a custom-made incubation setup, consisting of 200 mL borosilicate jacketed chambers maintained at controlled temperatures via a circulating bath and illuminated by four 10W LED light sources, all regulated through customized software. Magnetic stirrers ensured water homogenization during incubations.

Oxygen evolution was measured using optodes (Dipping Probe DP-PSt3, PreSens, Germany) connected to a fiber-optic oxygen meter (OXY4 SMA, PreSens, Germany), with data acquisition controlled via Measurement Studio 2 software (PreSens, Germany). To ensure accurate photosynthetic measurements and prevent underestimation due to oxygen oversaturation or carbon limitation, a biomass-to-seawater volume ratio of 0.03–0.05 g DW L^−1^ was maintained.

Leaf segments (∼3 cm ^2^) were initially incubated in darkness for 10 min to determine respiration (R) and then exposed to increasing PAR irradiance levels (0–801 µmol photons m⁻² s⁻¹) for 5 min at each intensity step. Light intensities were calibrated using a spherical (4π) quantum sensor (Biospherical Instruments, California, USA).

The maximum net photosynthetic rate (net-P_max_; µmol O_2_ g^-1^ DW h^-1^) was determined by averaging the maximum values above the saturating irradiance (E_k_ = net-P_max_/ α; µmol photon m^-2^ s^-1^). The gross maximum photosynthetic rate (gross-P_max_; µmol O_2_ g^-1^ DW h^-1^) was calculated as net-P_max_ + R. Photosynthetic efficiency (α) was calculated as the slope of the regression line fitted to the initial linear portion of the P vs. E curve. The compensation irradiance (E_c_; µmol photon m^-2^ s^-1^) was determined as the X-axis intercept of this linear portion (Fig. S4).

#### 2.4.2. Chlorophyll a fluorescence and absorptance

Chlorophyll *a* fluorescence from PSII was measured using a portable Diving-PAM fluorometer (Walz, Germany). Leaves were carefully cleaned of epiphytes and positioned in the DCL-8 leaf-clip holder to ensure a fixed distance between the tissue and the fiber optic sensor. Maximum quantum yield (F_v_/F_m_), basal fluorescence (F_₀_), and maximum fluorescence (F_m_) were recorded from plants kept in darkness overnight, while effective quantum yield (Ф_PSII_), F_₀_’, and F_m_’ were measured in light-acclimated plants exposed to actinic light (60 s, 247 µmol photon m⁻² s⁻¹). Dark and light adapted measurements were taken at the same leaf portion.

Absolute electron transport rate (ETR) was calculated using:

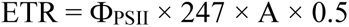

where A represents leaf absorptance (see below), 247 is the actinic light intensity, and 0.5 accounts for the assumption that half of the incident photons pass through PSII (Beer *et al*., 2014).

Leaf absorptance (A) was determined as:

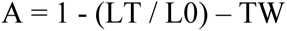

where LT is the transmitted light through the tissue, L0 is the total incident light emitted by the lamp, and TW is the transmitted light through a bleached leaf (2% bleach for 24 hours). Light transmission was measured using a miniature fiber optics quantum sensor (Diving-F1, Walz, Germany) integrated into the Diving-PAM fluorometer (Walz, Germany). The light sensor was secured in the leaf clip to maintain a consistent geometry between the leaf, lamp, and sensor.

Finally, non-photochemical quenching (NPQ) was calculated as:

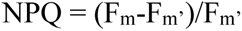

where F_m_ is the maximum fluorescence in dark-adapted plants, and F_m_’ is the fluorescence recorded under actinic light exposure.

#### 2.4.3. Pigment content

Leaf pigments were extracted from ∼15 mg fresh weight (FW) of homogenized leaf tissue using 100% acetone, with MgCO₃ added to prevent acidification (Dennison, 1990). Extracts were stored at 4°C in darkness for 24 hours. After centrifugation (1000 × g, 10 min), absorbance was measured at 470, 646, and 663 nm using a spectrophotometer and 1 mL cuvettes. Chlorophyll *a* (Chl *a*), chlorophyll *b* (Chl *b*), and total carotenoid concentrations were calculated using the equations of Lichtenthaler & Wellburn (1983) and expressed as mg g⁻¹ FW.

#### 2.4.4. Total phenolic content, antioxidant capacity and lipid peroxidation

Phenolic and antioxidant compounds were extracted from dried, ground leaf tissue (0.02 g DW) using 1 mL of 80% methanol. Samples were incubated in darkness for 24 hours and then centrifuged at 10,000 rpm for 10 minutes. The resulting extracts were used to quantify total phenolic content and antioxidant capacity.

Total phenolic content was determined using a modified Folin–Ciocalteu assay with gallic acid as the standard (Singleton & Rossi, 1965). A 0.01 mL aliquot of the methanolic extract was diluted in 1 mL of distilled water (dH₂O), followed by the addition of 0.1 mL Folin–Ciocalteu reagent and 0.3 mL Na_2_CO₃-saturated dH₂O. The mixture was homogenized, heated at 40°C for 3 minutes, and absorbance was measured at 765 nm using a spectrophotometer. Total phenolic content was expressed as gallic acid equivalents (mg Eq. GA g⁻¹ DW).

Radical scavenging activity was quantified from the same methanolic extracts following Sabeena-Farvin & Jacobsen (2013). The reaction mixture was prepared by mixing 0.1 mL of a 1:10 diluted extract (in 80% methanol) with 1 mL of 60 μM 2,2-diphenyl-1-picrylhydrazyl (DPPH) dissolved in 90% methanol. Absorbance was recorded at 517 nm after 30 minutes of DPPH addition. Total antioxidant capacity was expressed as ascorbic acid equivalents (mg Eq. AA g⁻¹ DW).

Oxidative damage was determined by quantifying malondialdehyde (MDA). Lipid peroxidation was quantified using the thiobarbituric acid-reactive substances (TBARS) assay, following Hodges *et al*. (1999) and Correia *et al*. (2006). Frozen leaf tissue (0.2 g FW) was subjected to mechanical grinding in 2 mL of 80% ethanol. The homogenate was centrifuged (10 min, 3000 × g, 4°C), and the supernatant was mixed with 20% trichloroacetic acid (TCA) and 0.5% thiobarbituric acid (TBA). Blanks were prepared by mixing the supernatant with 20% TCA. Samples were incubated at 90°C for 30 minutes and centrifuged again (3000 × g, 10 min). Absorbance of the extracted supernatants was measured at 440, 532, and 600 nm using a spectrophotometer. Lipid peroxidation was expressed as malondialdehyde equivalents (Eq. MDA) using a molar extinction coefficient of 155 mM⁻¹ cm⁻¹, according to Hodges *et al*. (1999).

#### 2.4.5. Nitrogen uptake and nitrate reductase activity

Nitrogen uptake was quantified by incubating shoots in artificial seawater enriched with 5 μM of isotopically labeled nitrate (K¹⁵NO₃, 99 atom % ¹⁵N; Cambridge Isotope Laboratories) for 30 minutes. This concentration is similar to nitrate levels observed in surfgrass meadows during upwelling events, which tend to be most intense and frequent in the region throughout the spring and summer (Espinosa-Carreón *et al*., 2001). Incubations were conducted in separate plastic bags (2 L, n = 3), which were sealed and left floating during the incubation period inside the experimental tanks, ensuring that temperature and irradiance matched those of the respective treatment. Each incubation encompassed eight randomly selected shoots per tank and treatment, maintaining a weight-to-volume ratio of ∼4.5 g FW L⁻¹. To preserve vegetative integrity, a small rhizome segment (∼1 cm) was retained in each shoot. Nitrogen uptake rates were calculated using the equations of Sandoval-Gil *et al*. (2016). Dried and ground leaf tissues were analyzed for ¹⁵N enrichment and total nitrogen content (N-content) at the University of California Davis-Stable Isotope Facility, using an elemental analyzer interfaced with a continuous-flow isotope ratio mass spectrometer (IRMS). To assess nitrogen assimilation, nitrate reductase (NRA) activity was measured *in vivo* from leaf segments of three shoots per tank and treatment, following Alexandre *et al*. (2004), using leaves distinct from those analyzed for nitrate uptake.

#### 2.4.6. Non-structural carbohydrates

Total soluble non-structural carbohydrates (NSCs), including free sugars and starch, were quantified using the colorimetric phenol-sulfuric acid method (Dubois *et al*., 1956), with glucose as the standard. Leaf tissues were oven-dried at 60°C to a constant weight, then ground into a fine powder. The powdered tissue was digested in 0.1 N HCl (60°C, 3 h), centrifuged (4000 × g, 5 min), and the supernatant was mixed with 3% phenol and concentrated sulfuric acid. Absorbance was recorded at 490 nm using a spectrophotometer. NSC content was expressed as mg g⁻¹ DW.

#### 2.4.7. Daily Productivity

Daily productivity (DP; µmol O₂ g⁻¹ DW day⁻¹) was estimated by integrating the P vs. E response curves with the averaged diurnal irradiance cycle recorded in the experimental system. Unlike methods that rely on fixed photosynthetic efficiency (α) and net-P_max_, we applied polynomial functions (3rd or 4th degree) fitted to the P vs. E data from each experimental unit (EU, n = 4), with the y-intercept fixed at R.

Diurnal irradiance was recorded using Onset HOBO MX2202 sensors (n = 3) placed at mid-water depth to capture both direct and reflected sunlight within the aquaria. Light intensity (Lux) was converted to photons m^-2^ s^-1^ using a conversion factor of 0.0185. Each recorded irradiance value was entered into the polynomial equation to estimate instantaneous photosynthetic rates. Integrating these rates over the whole daylight period yielded cumulative daily productivity.

#### 2.4.8. Relative leaf growth rate

All leaves of each shoot were marked just above the sheath at the beginning of both the warming and post-warming phases using the hole punch method adapted from Zieman (1974). At the end of each experimental phase, all marked shoots were harvested, and newly formed leaf tissue (i.e., below the mark) was dried and weighed for each shoot. Shoot growth was expressed as relative leaf growth rate (RGR; mg g⁻¹ DW day⁻¹), calculated with respect to total shoot biomass.

### 2.5. Statistical analysis

#### 2.5.1. Univariate analysis

All statistical analyses were conducted in R (R Core Team, 2020), with a significance level of α = 0.05. Prior to hypothesis testing, data normality was assessed using the Shapiro-Wilk test, and homogeneity of variances was evaluated using Bartlett’s test (package car). Variables that deviated from normality were log-transformed to improve distribution symmetry (e.g., chl *b* content and nitrate uptake rate). A two-way ANOVA was performed to assess the main and interactive effects of treatment (three levels: control, severe-MHW and extreme-MHW) and experimental phase (two levels: warming phase and post-warming phase) on all measured physiological descriptors. A standard type III ANOVA was used for the variables that met the parametric assumptions. For heteroscedastic variables (e.g., F_m_, and NRA activity), ANOVA models were adjusted using White’s correction to account for variance heterogeneity. Tukey’s HSD post-hoc test was applied when significant differences were detected to compare means among treatments and time points. The non-parametric Games-Howell test was used for *post-hoc* comparisons in heteroscedastic variables.

#### 2.5.2. Multivariate analysis

Principal Component Analysis (PCA) was performed using the *prcomp* function in R on a correlation matrix (scale. = TRUE). The number of components retained was based on the Kaiser criterion (eigenvalue > 1). Loadings were used to identify key variables contributing to each axis, and scores were used to visualize sample distribution across treatments and time points. To reduce redundancy and improve the interpretability of the PCA, a variable selection approach was applied prior to the analysis. Firstly, variables that were mathematically derived from others were excluded to prevent artificial correlations and overrepresentation of specific physiological processes. Secondly, variables that did not exhibit significant responses to treatments in the univariate analysis were removed. Lastly, correlated variables that reflected similar but independently measured physiological processes were combined into composite indices (Table S3). Variables were standardized (z-score), and grouped indices were averaged prior to analysis.

Multivariate differences were also tested using Permutational ANOVA (PERMANOVA, *adonis2* function, *vegan* package) based on a Euclidean distance matrix. Treatment, experimental Phase, and their interaction were included as fixed factors, with 999 permutations. Pairwise comparisons among Treatment × Experimental Phase combinations were also performed.

## 3. Results

### 3.1. Photosynthesis and respiration

Photosynthetic capacity varied across treatments and experimental phases, with significant treatment effects detected for net-P_max_, E_k_, E_c_ and R (Table 1). The most pronounced changes occurred under the extreme-MHW treatment, where net-Pₘₐₓ and Eₖ decreased by 12% and 22%, respectively, during the warming phase, and by 39% and 19% during the post-warming phase, relative to the control plants (Figs. 2A, C). These reductions were accompanied by significant increases in R, which rose by 77% during the warming phase and by 113% post-warming (Fig. 2F). Similarly, E_c_ increased by 53% during warming and by 181% during post-warming under extreme-MHW conditions (Fig. 2E). In contrast, no significant treatment effects were detected for gross-P_max_ or α (Figs. 2B, D).

**Figure 2.**
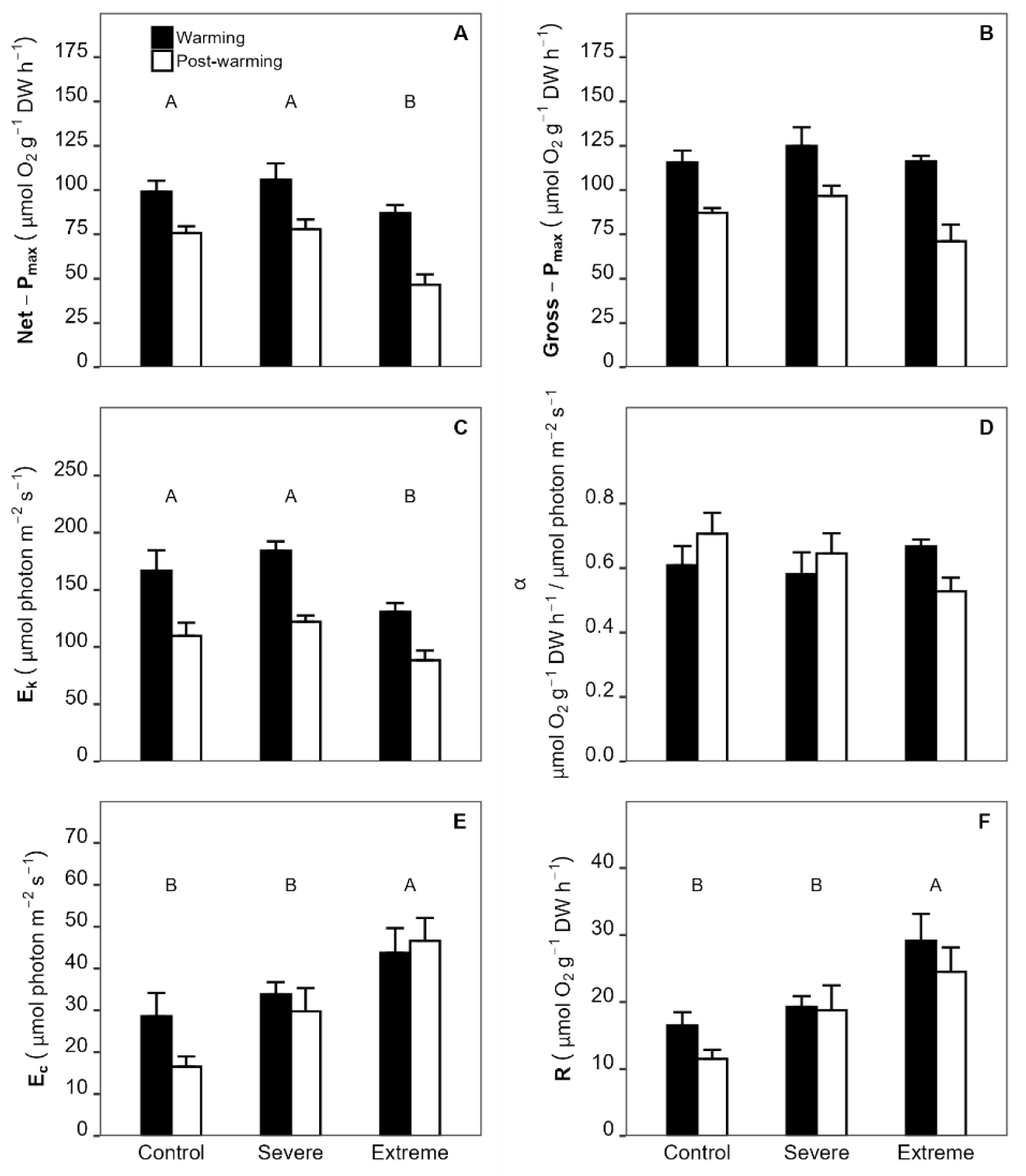
Photosynthesis–irradiance curve derived measured in *Phyllospadix scouleri* under three temperature treatments: Control (18.5 ± 1.5°C), Severe-MHW (23.5 ± 1.5°C) and Extreme-MHW (26.5 ± 1.5°C). Measurements were taken at the end of the warming and post-warming phases. Bars represent mean ± SE (n = 4). Two-way ANOVA results for treatment, experimental phase, and their interaction are shown. Significant differences among treatments (p < 0.05, Tukey HSD) are indicated by different uppercase letters when the main effect of treatment was significant. See Table 1 for full statistical results. Variables: (A) Net maximum photosynthetic rate (Net-P_max_), (B) Gross-P_max_, (C) Saturation irradiance (E_k_), (D) Photosynthetic efficiency (α), (E) Compensation irradiance (E_c_), (F) Respiration rate (R).

**Table 1.**
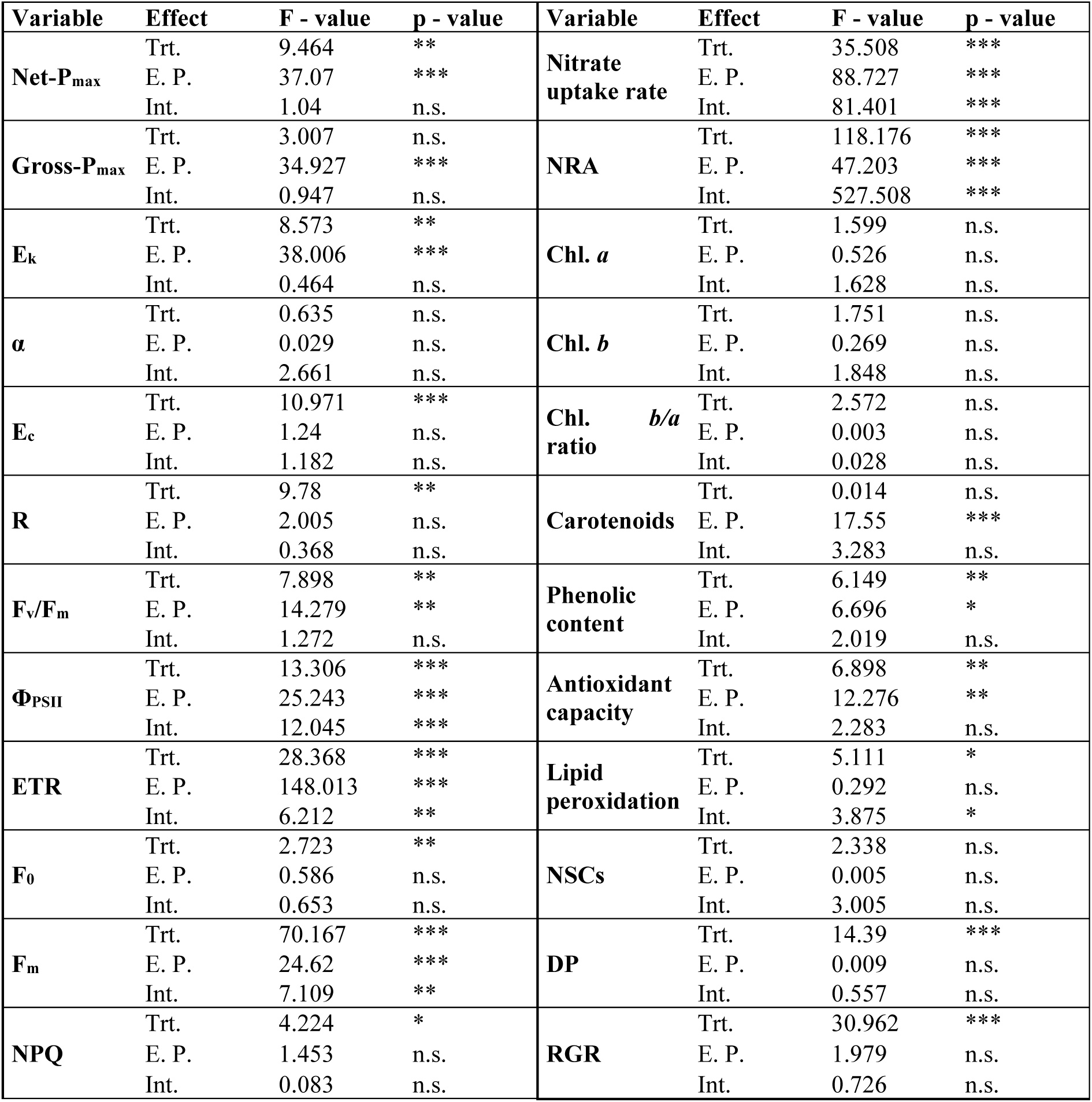
Results of two-way ANOVA testing the effects of treatment (Trt., 3 levels: Control, HW-1, HW-2), experimental phase (E.P., 2 levels: T-1 and T-2), and their interaction (Int.) on all biological descriptors. Significance codes: *p < 0.05; **p < 0.01; ***p < 0.001, n.s. p > 0.05.

### 3.2. Chlorophyll *a* fluorescence and absorptance

The photochemical performance of photosystem II was significantly affected by warming, as evidenced by a decline in the maximum photochemical efficiency (F_v_/F_m_) in plants exposed to the extreme-MHW treatment (Table 1). Compared to the control, F_v_/F_m_ decreased by 3% during the warming phase and by 7% in the post-warming phase (Fig. 3A).

**Figure 3.**
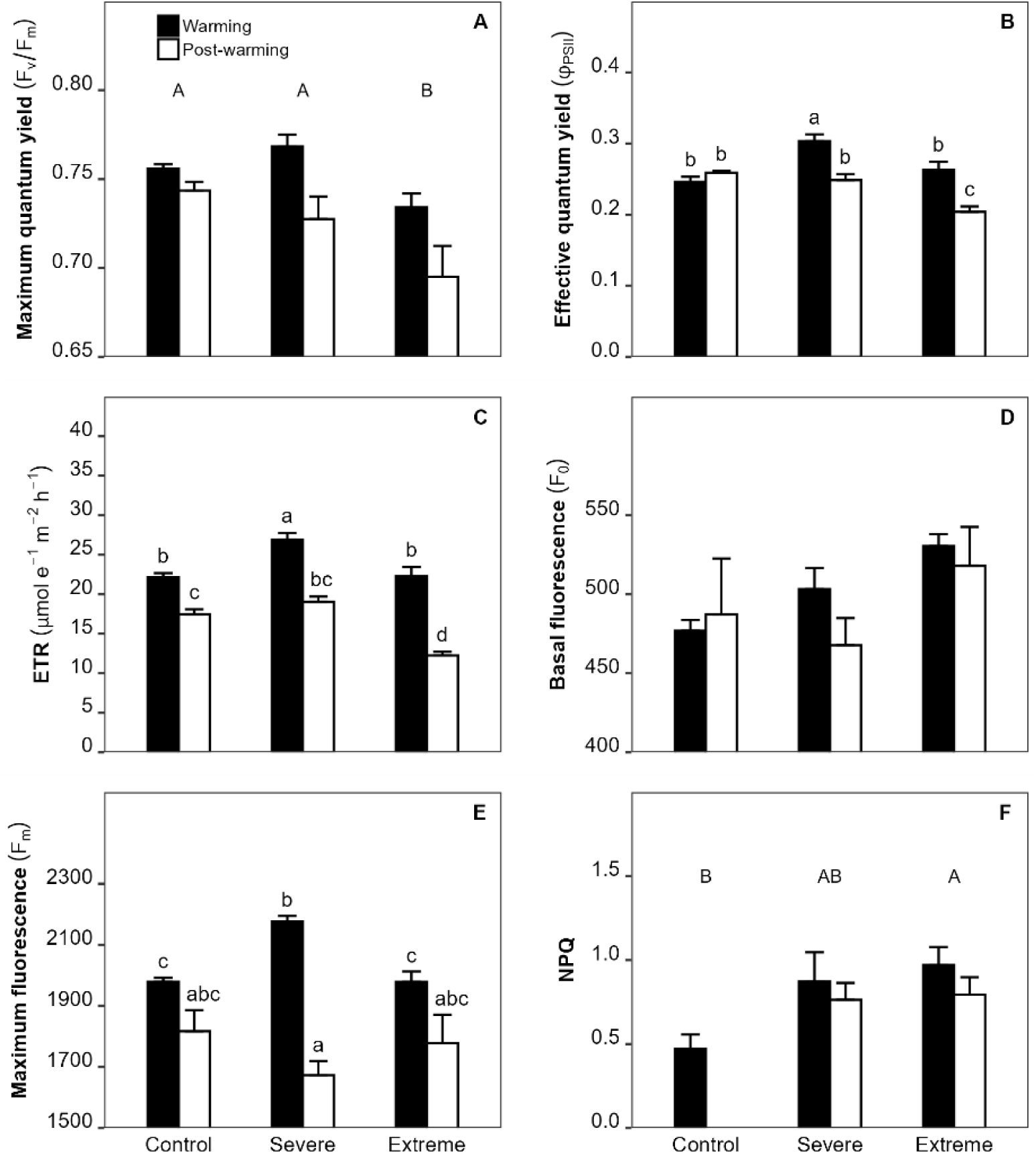
Photochemistry parameters measured in *Phyllospadix scouleri* under three temperature treatments: Control (18.5 ± 1.5°C), Severe-MHW (23.5 ± 1.5°C) and Extreme-MHW (26.5 ± 1.5°C). Measurements were taken at the end of the warming and post-warming phases. Bars represent mean ± SE (n = 4). Two-way ANOVA results for treatment, experimental phase, and their interaction are shown. Significant differences among treatments (Tukey HSD, p < 0.05) are indicated with uppercase letters when the main effect of treatment was significant, and with lowercase letters for pairwise comparisons among treatments within each phase when a significant interaction was detected. See Table 1 for full statistical results. Variables: (A) Maximum quantum yield (F_v_/F_m_), (B) Effective quantum yield (Φ_PSII_), (C) Electron transport rate (ETR), (D) Basal fluorescence (F_0_), (E) Maximum fluorescence (F_m_), (F) Non-photochemical quenching (NPQ).

The effective quantum yield (Φ_PSII_) showed a significant interaction between treatments and experimental phase (Table 1). The highest Φ_PSII_ was observed in plants subjected to the severe-MHW treatment during the warming phase, significantly exceeding values in both the control and extreme-MHW groups. By the post-warming phase, Φ_PSII_ had declined in both MHW treatments, with the extreme-MHW group exhibiting the steepest drop, 21% lower than the control (Fig. 3B).

The electron transport rate (ETR) followed a pattern similar to that of Φ_PSII_, showing a significant interaction between treatment and experimental phase (Table 1). During the warming phase, ETR was highest in the severe-MHW treatment, approximately 21% higher than the control. In contrast, ETR declined sharply in the extreme-MHW group during the post-warming phase, reaching values 30% lower than the control (Fig. 3C).

Basal fluorescence (F_0_) was not significantly affected by experimental phase, treatment, or their interaction (Table 1). However, an increasing trend was observed during the warming phase across both MHW groups (Fig. 3D). By the post-warming phase, F_0_ values remained higher only in the extreme-MHW treatment.

In contrast, maximum fluorescence (F_m_) was significantly influenced by experimental phase, treatment, and their interaction (Table 1). During the warming phase, plants in the severe-MHW treatment exhibited the highest F_m_ values. By the post-warming phase, F_m_ declined in all treatments (Fig. 3E).

Non-photochemical quenching was lowest in control plants during the warming phase (Fig. 3F). Due to inconsistencies in measurements, control values during the post-warming phase were excluded from the analysis. Both MHW treatments showed increased NPQ levels, but only the extreme-MHW group was significantly higher than the control (Table 1). NPQ peaked during the warming phase in this treatment, reaching values 106% higher than the control.

### 3.3. Pigment content

Pigments contents were not significantly affected by the experimental treatments. However, for carotenoids, pigment levels varied between experimental phases (Tables 1, 2). Plants in the severe-MHW treatment showed the highest carotenoid concentrations during the warming phase, while those in the extreme-MHW treatment reached their peak values during the post-warming phase. These differences, however, were not statistically significant.

**Table 2.** Mean (± SE) values of pigment concentration (chlorophyll *a*, chlorophyll *b*, Chl *b/a* ratio, carotenoids), phenolic content, antioxidant capacity, and lipid peroxidation across treatments (Control, Severe, Extreme) and experimental phases (Warming and Post-warming). Letters in parentheses indicate statistically significant differences based on post hoc comparisons (*p* < 0.05). Uppercase letters denote differences among treatments; lowercase letters indicate interaction effects.

| Variable | Treatment | Experimental phase |  |
| --- | --- | --- | --- |
|  |  | Warming | Post-warming |
| Chlorophyll *a* | Control | 1.1 ± 0.07 | 1.02 ± 0.04 |
| | Severe | 1.37 ± 0.13 | 1.12 ± 0.18 |
| | Extreme | 1.11 ± 0.1 | 1.24 ± 0.03 |
| Chlorophyll *b* | Control | 0.49 ± 0.03 | 0.47 ± 0.01 |
| | Severe | 0.55 ± 0.03 | 0.47 ± 0.08 |
| | Extreme | 0.52 ± 0.04 | 0.59 ± 0.02 |
| Chl. *b/a* ratio | Control | 0.44 ± 0.01 | 0.45 ± 0.01 |
| | Severe | 0.41 ± 0.01 | 0.41 ± 0.01 |
| | Extreme | 0.45 ± 0.03 | 0.44 ± 0.01 |
| Carotenoids | Control | 0.27 ± 0 | 0.27 ± 0.01 |
| | Severe | 0.31 ± 0.02 | 0.26 ± 0.03 |
| | Extreme | 0.28 ± 0.03 | 0.35 ± 0 |
| Phenolic content | Control (A) | 5.9 ± 0.35 | 6.12 ± 0.63 |
| | Severe (B) | 4.3 ± 0.13 | 5.82 ± 0.34 |
| | Extreme (B) | 4.66 ± 0.24 | 5.1 ± 0.1 |
| Antioxidant capacity | Control (A) | 2.77 ± 0.49 | 3.15 ± 0.29 |
| | Severe (AB) | 1.69 ± 0.06 | 3.11 ± 0.2 |
| | Extreme (B) | 1.72 ± 0.21 | 2.22 ± 0.14 |
| Lipid peroxidation | Control | 29.86 ± 1.24 (a) | 29.58 ± 1.98 (a) |
| | Severe | 20.75 ± 1.85 (b) | 27.71 ± 1.72 (a) |
| | Extreme | 31.86 ± 2.02 (a) | 27.84 ± 2.86 (a) |

### 3.4. Total phenolic content, antioxidant activity and lipid peroxidation

Leaf phenolic content was significantly reduced in plants exposed to warming (Table 1). Plants in the severe-MHW and extreme-MHW treatments exhibited reductions of approximately 27% and 21%, respectively, during the warming phase compared to the control (Table 2). By the post-warming phase, the phenolic content increased in both treatments but remained 5% lower in severe-MHW and 17% lower in extreme-MHW plants compared to control levels.

Total antioxidant capacity was also significantly affected by treatment (Table 1). During the warming phase, both severe-MHW and extreme-MHW plants exhibited approximately 40% lower antioxidant capacity than control plants (Table 2). However, only the extreme-MHW group differed significantly from the control. By the post-warming phase, severe-MHW plants recovered to control levels, while values in the extreme-MHW group remained 30% lower than those of the control.

Lipid peroxidation was significantly affected by treatment, with a significant interaction with experimental phase (Table 1). In the severe-MHW treatment, lipid peroxidation showed a significant decrease during the warming phase, reaching values 31% lower than the control, but returned to near-control levels by the post-warming phase (Table 2). In contrast, lipid peroxidation in extreme-MHW plants did not vary significantly.

### 3.5. Nitrogen uptake and NRA

Nitrate uptake was significantly affected by the interaction between treatment and experimental phase (Table 1). During the warming phase, plants in the extreme-MHW treatment exhibited the highest uptake rates, while in the severe-MHW group uptake was 38% lower than in the control (Fig. 4A). By the post-warming phase, severe-MHW and extreme-MHW plants exhibited contrasting responses, but both had showed higher uptake rates than the control, increasing by 82% and 43%, respectively.

**Figure 4.**
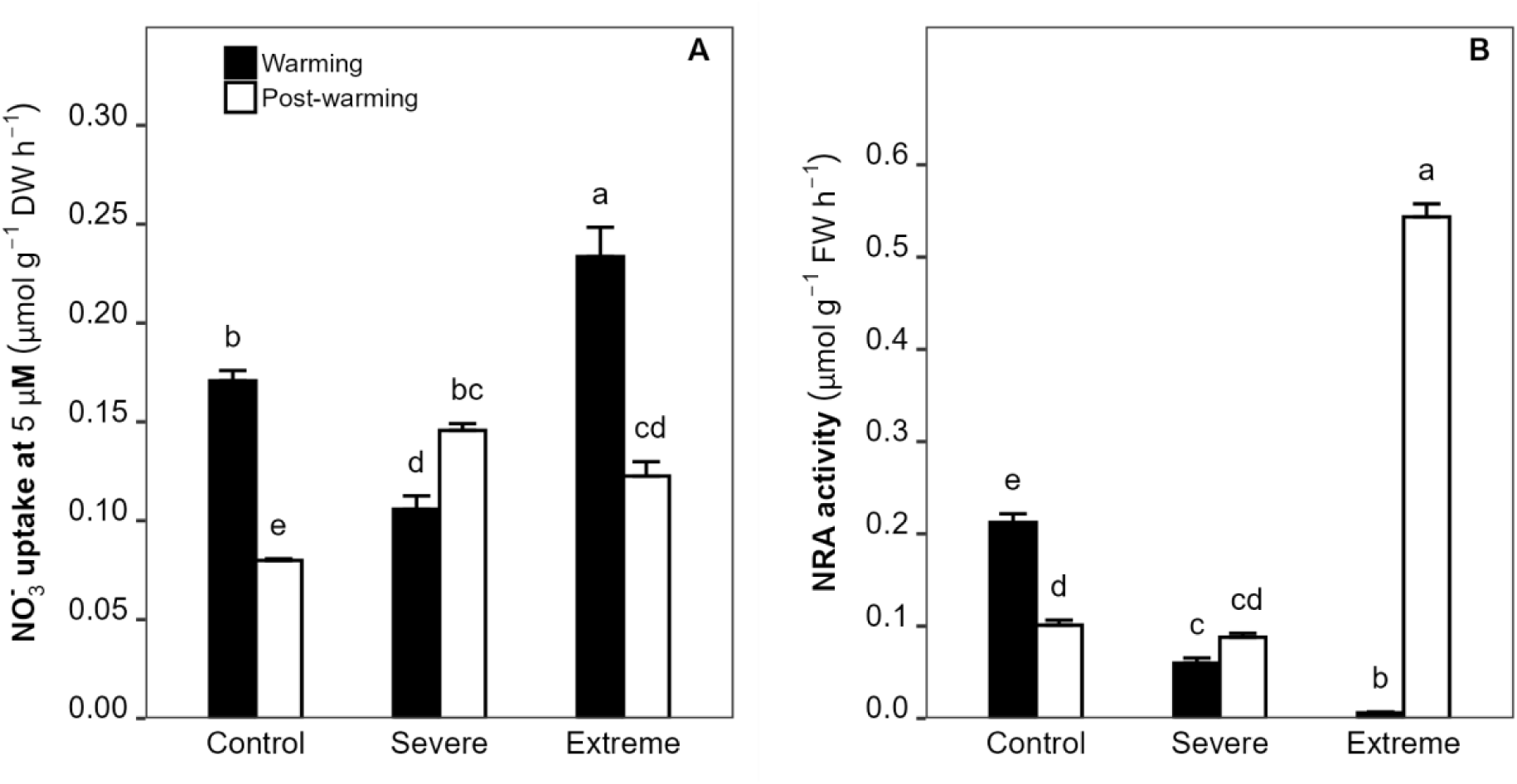
Nitrogen metabolism in *Phyllospadix scouleri* under three temperature treatments: Control (18.5 ± 1.5 °C), Severe-MHW (23.5 ± 1.5 °C), and Extreme-MHW (26.5 ± 1.5 °C). Measurements were taken at the end of the warming and post-warming phases. Bars represent mean ± SE (n = 4). Two-way ANOVA results for treatment, experimental phase, and their interaction are shown. When a significant interaction was detected, lowercase letters indicate significant pairwise differences among treatments within each phase (Tukey HSD, p < 0.05). See Table 1 for full statistical results. Variables: (A) Nitrate uptake rate at 5 μM, (B) Nitrate reductase activity (NRA).

Nitrate reductase activity (NRA) was also significantly affected by the interaction between treatment and experimental phase (Table 1). During the warming phase, control plants showed a modest 16% reduction in NRA compared to initial conditions. In comparison, severe-MHW and extreme-MHW plants exhibited sharp declines of 71% and 97%, respectively, relative to the control (Fig. 4B). By the post-warming phase, NRA continued to decline in control plants, remained stable in the severe-MHW group, and showed a drastic increase in the extreme-MHW treatment, reaching values 438% higher than those of the control.

### 3.6. Carbohydrates, productivity and growth

Leaf non-structural carbohydrate (NSC) content did not vary significantly across treatments (Table 1), although plants in the severe-MHW treatment exhibited a 52% increase in NSC content relative to the control (Fig. 5A).

**Figure 5.**
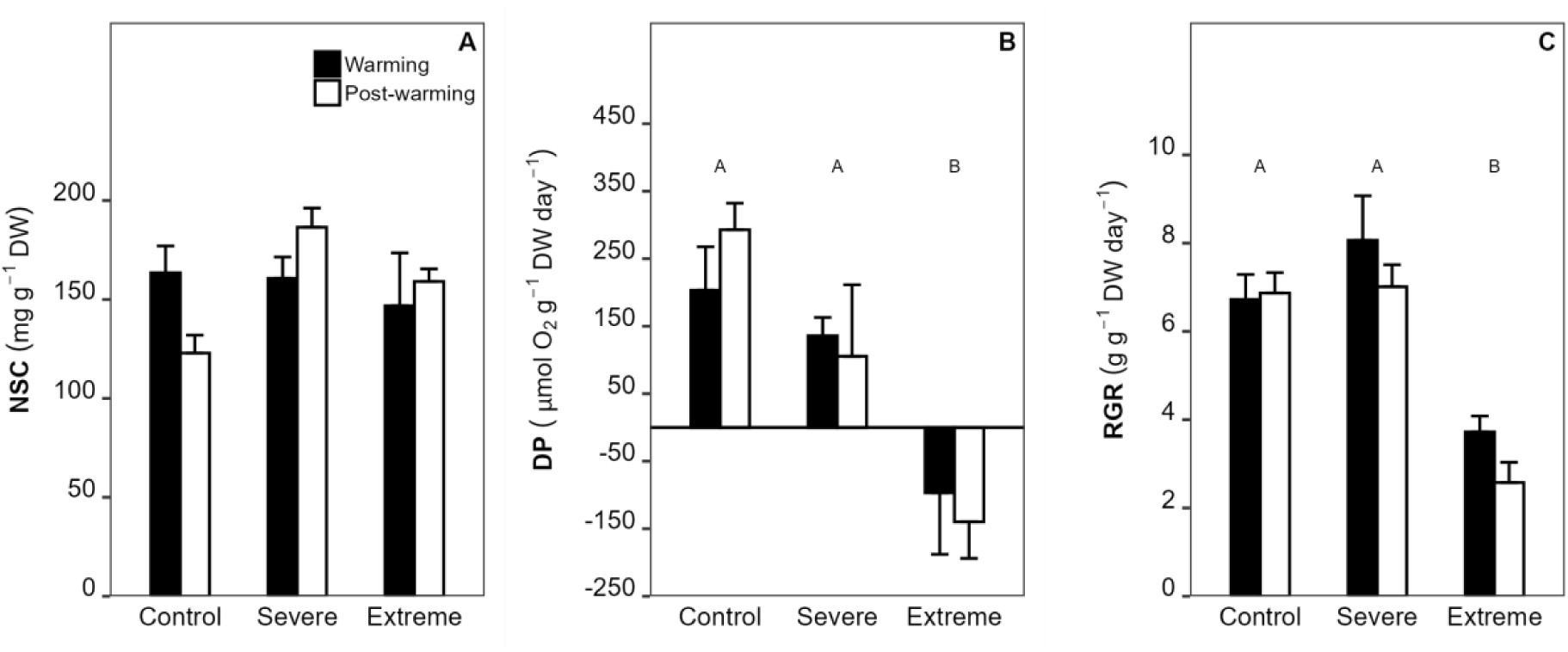
Energy and growth metrics measured in *P. scouleri* under three temperature treatments: Control (18.5 ± 1.5°C), Severe-MHW (23.5 ± 1.5°C) and Extreme-MHW (26.5 ± 1.5°C). Measurements were taken at the end of the warming and post-warming phases. Bars represent mean ± SE (n = 4). Two-way ANOVA results for treatment, experimental phase, and their interaction are shown. Significant differences among treatments (p < 0.05, Tukey HSD) are indicated by different uppercase letters when the main effect of treatment was significant. See Table 1 for full statistical results. Variables: (A) Non-structural carbohydrates (NSC), (B) Daily productivity (DP), (C) Relative growth rate (RGR).

Daily productivity (DP) was significantly influenced by treatment (Fig. 5B). Control plants experienced a substantial 58% decline in DP during the warming phase compared to initial conditions. Plants in the severe-MHW treatment showed slightly lower DP than the control, although the difference was not statistically significant. In contrast, the extreme-MHW treatment resulted in significantly lower DP values compared to both the control and severe-MHW, with negative productivity observed during both the warming and post-warming phases.

Leaf relative growth rate (RGR) was strongly affected by treatment (Fig. 5C). control and severe-MHW plants maintained similar RGR values throughout the experiment. In contrast, plants exposed to the extreme-MHW treatment showed a 45% reduction during the warming phase and a 62% reduction during the post-warming phase compared to the control.

### 3.7. Multivariate results

Principal Component Analysis (PCA) revealed clear patterns of multivariate physiological variation across treatments and experimental phases (Fig. 6). The first three principal components explained 78.2% of the total variance, with PC1 (31.8%) and PC2 (24.4%) accounting for the greatest proportion (Table S4). Samples from the extreme-MHW treatment, particularly during the post-warming phase, were clearly separated from those of the control and severe-MHW groups along PC1. This separation was driven by decreases in respiration, nitrogen assimilation, and carotenoids, whereas control and severe-MHW samples expressed higher values of leaf growth and photosynthesis. PC2 further distinguished extreme-MHW samples during the warming phase, which were associated with increased nitrate uptake and respiration. PC3 (22.0%) captured additional variation related to antioxidant capacity and oxidative stress, contributing to the separation of samples within the control and severe-MHW treatments during the post-warming phase.

**Figure 6.**
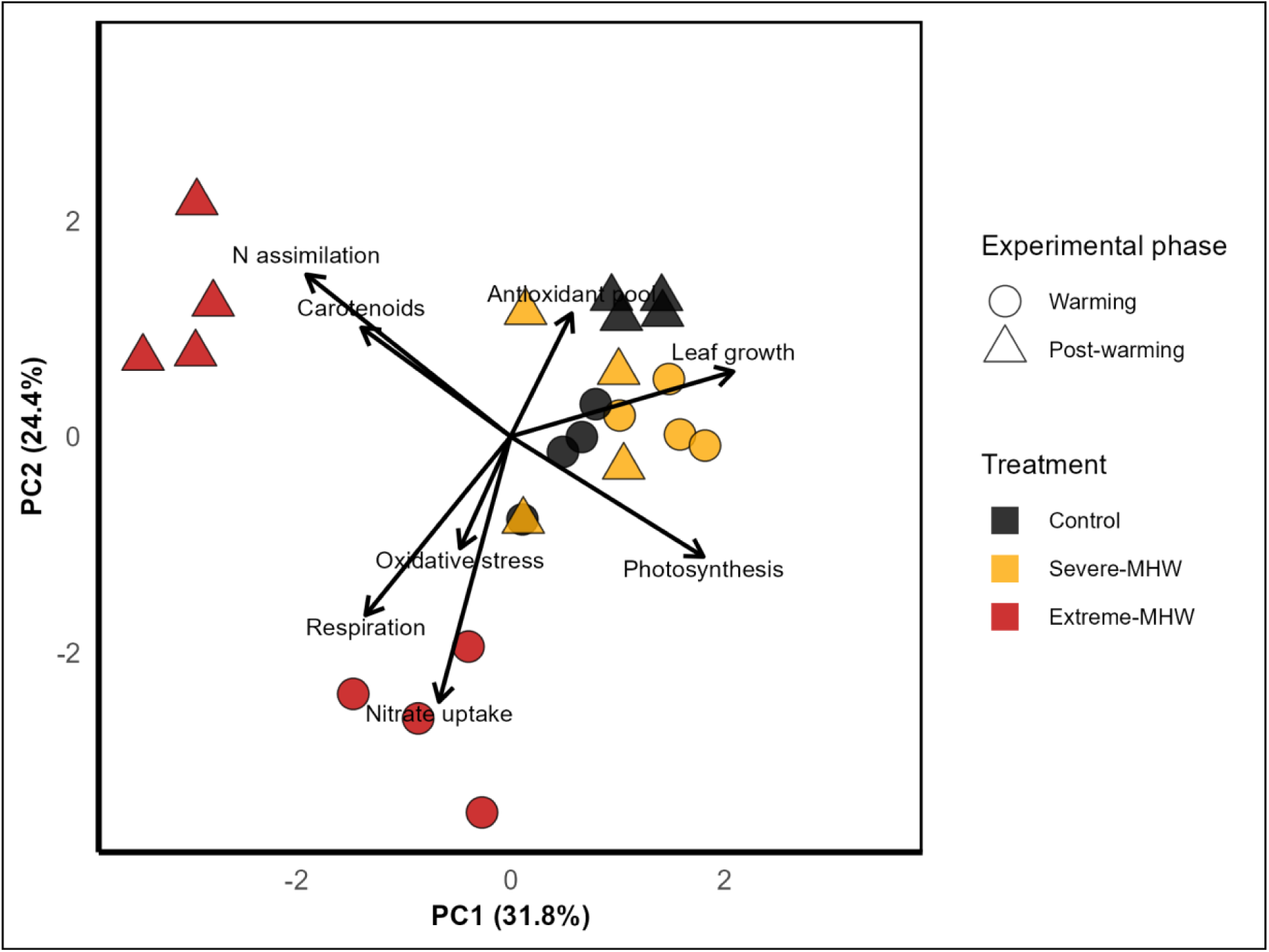
Principal Component Analysis (PCA) performed with the physiological responses of *Phyllospadix scouleri* to three temperature treatments: Control (18.5 ± 1.5°C), Severe-MHW (23.5 ± 1.5°C) and Extreme-MHW (26.5 ± 1.5°C). Measurements were taken at the end of the warming and post-warming phases. The first two principal components explain 56.2% of the total variance (PC1: 31.8%, PC2: 24.4%). Arrows represent loadings of key physiological variables contributing to each axis. See Table S4 for further PCA details.

PERMANOVA confirmed significant effects of treatment (R^2^ = 0.33, P = 0.001), experimental phase (R^2^ = 0.15, P = 0.001), and their interaction (R^2^ = 0.23, P = 0.001) on multivariate physiological profiles (Table S5). Pairwise comparisons revealed that the extreme-MHW treatment differed significantly from both the control and severe-MHW treatments during both the warming and post-warming phases, consistent with the separation observed in the PCA. The severe-MHW group differed from the control during the warming phase but not during the post-warming phase, suggesting partial recovery or convergence toward baseline conditions by the end of the experiment.

## 4. Discussion

As marine heatwaves (MHWs) intensify under climate change, foundation species such as seagrasses are being pushed toward their physiological limits, threatening the resilience of the coastal ecosystems they support. This study examines how the surfgrass *Phyllospadix scouleri* responds physiologically to increasing thermal stress, revealing the upper limits of its thermal tolerance (Fig. 7). While *P. scouleri* maintained physiological stability under a severe MHW (T_mean_ = 23.5 °C, T_max_ = 25 °C), exposure to an extreme MHW (T_mean_ = 26.5°C, T_max_ = 28 °C) caused pronounced photosynthetic impairment, supression of nitrogen assimilation, elevated respiration and a negative carbon balance, ultimately leading to reduced leaf growth. These findings indicate that future, more intense MHWs could exceed the thermal tolerance of surfgrass populations, potentially threatening their ecological functioning and long-term persistence.

### 4.1. Mechanisms of Physiological Tolerance vs. Breakdown

Exposure to a severe MHW did not negatively affect the photosynthetic performance of *P. scouleri*. Both net- and gross-P_max_ remained stable, and the functionality of the photosynthetic apparatus at thylakoid level (i.e., ETR and Φ_PSII_) was even enhanced, indicating that the species’ thermal tolerance (at least at photosynthetic metabolism level) was not surpassed. Moderate warming has also been shown to increase photosynthesis in *P. torreyi* (Vivanco-Bercovich *et al*., 2022) and other species (Beca-Carretero *et al*., 2018; Lawrence & Bolton, 2022), likely due to enhanced enzyme activity and protein mobility in the thylakoid membrane.

In contrast, exposure to the extreme MHW appeared to push *P. scouleri* beyond a physiological threshold, resulting in marked declines in its photosynthetic performance, including reductions in net-P_max_, gross-P_max_, E_k_, F_v_/F_m_, Φ_PSII_, and ETR. Photosynthetic alterations in seagrasses under heat stress are well documented (e.g., Costa *et al*., 2021; Nguyen *et al*., 2021; Deguette *et al*., 2022), and are often linked to damage in key components of PSII, such as light-harvesting complexes, reaction centers, and oxygen-evolving complexes (Mathur *et al*., 2014). As chlorophyll *a* and *b* levels remained stable, it is likely that functional damage to thylakoid membranes was the main cause of photosynthetic inhibition (Marín-Guirao *et al*., 2018). The observed decrease in F_v_/F_m_, along with increased basal fluorescence (F_0_), further supports stress to PSII structure (Allakhverdiev *et al*., 2008).

Our results showed that non-photochemical quenching (NPQ) increased in *P. scouleri* exposed to the extreme MHW during both experimental phases. This enhancement reflects the activation of a photoprotective mechanism that dissipates excess light energy as heat through the xanthophyll cycle, thereby preventing de production of reactive oxygen species and oxidative stress (Ruocco *et al*., 2019; Murchie & Ruban, 2020; Vivanco-Bercovich *et al*., 2024). Notably, no increase in lipid peroxidation was detected, suggesting either absence of oxidative damage to membranes or sufficient antioxidant capacity to counteract stress (Gururani *et al*., 2015). Likewise, total antioxidant capacity and phenolic content remained unchanged, indicating marginal stimulation of antioxidant defenses under warm conditions.

Warming also affected nitrogen metabolism in *P. scouleri*, but responses varied with MHW intensity. As temperature increased NRA declined, consistent with earlier reports (Bonet-Melià *et al*., 2023). However, contrary to expectations, nitrate uptake decreased under severe warming but increased under extreme conditions. This pattern may reflect complex interactions among nitrogen incorporation, assimilation, translocation, and storage investment and mobilization processes (Touchette & Burkholder, 2007; Alexandre *et al*., 2010), though these mechanisms remain to be fully explored. Under extreme MHW, high uptake rates coupled with low assimilation suggest a compensatory strategy to maintain internal nitrogen stores, as documented in seaweeds (Gordillo, 2012). Nevertheless, these disruptions were transient, with both uptake and assimilation recovering, or even surpassing, control levels once warming ceased.

These results demonstrate that when MHWs exceed a critical threshold, *P. scouleri* undergoes a disruption in carbon balance atributable to a mismatch between photosynthetic production and respiratory demand. While moderate warming (∼25–26 °C) was tolerated without excessive respiratory demand (Vivanco-Bercovich *et al*., 2022; 2024), exposure to the extreme MHW caused sustained elevation in R. Given the temperature sensitivity of seagrass R (Marín-Guirao *et al*., 2016; Beca-Carretero *et al*., 2018), it is probable that this pushed *P. scouleri* beyond a metabolic tipping point, leading to negative daily productivity. The observation that the NSC content of leaves remained stable suggests that plants relied on alternative carbon pools, most likely rhizome starch reserves, reflecting the flexible and spatially complex carbon economy of seagrasses (Marín-Guirao *et al*., 2018; Ruocco *et al*., 2021). Reduced leaf growth under extreme conditions may thus reflect an energy conservation strategy, prioritizing cellular maintenance and repair over biomass accumulation (Ruocco *et al*., 2019), a potential early indicator of long-term population decline.

This shift from tolerance to functional collapse was clearly captured in the multivariate analyses. The results of the PCA analysis revealed that *P. scouleri* plants exposed to the severe-MHW recovered to near-control physiological states following the cessation of warming (i.e. post-warming phase). At the same phase, plants subjected to the extreme-MHW remained clearly separated, driven by sustained R, impaired photosynthetic performance, reduced nitrate assimilation, and limited growth. These differences were further corroborated by the PERMANOVA. Respiration emerged as a key variable contributing to this divergence, illustrating how sustained energetic imbalance can cascade across physiological pathways and limit recovery capacity. Together, these findings highlight that while *P. scouleri* can recover from moderate thermal stress, exposure to extreme warming events may induce prolonged dysfunction that compromises its resilience.

### 4.2. Implications for Future Climate Change and Conclusions

Previous research has shown that exposure to 25°C can induce physiological alterations in *P. scouleri* without excerting a significant effect on productivity and growth (Vivanco-Bercovich *et al*., 2024). However, in this study, the extreme MHW scenario (T_mean_ = 26.5°C; T_max_ = 28°C) caused severe photosynthetic impairment, carbon imbalance, and growth suppression, suggesting a critical thermal threshold beyond which survival may be compromised. Taken together, the available evidence suggests that 26.5°C (as a daily average for at least seven days) may represent a tipping point (T_lim_) for surfgrass thermal resistance in the northern Baja California Peninsula.

This threshold gains further significance when considered in the context of recent thermal trends in the region. The mean maximum summer temperature (T_max_) in the study region was ∼20.5°C between 1990 and 2019, rising to 21.5°C in the period 2014–2019. This suggests that the current thermal safety margin (TSM = T_max_ - T_lim_) for *P. scouleri* populations in northern Baja California Peninsula is approximately 5 - 6°C. While this study focused on a local population, the physiological limit observed here aligns closely with the ∼27.5 °C upper thermal limit reported for temperate seagrasses worldwide (Marbà *et al*., 2022), reinforcing its relevance for regional thermal vulnerability assessments.

Continued ocean warming and increasingly intense MHWs may significantly reduce this margin in the near future. Recent temperature records indicate that sea surface temperature in the study region has repeatedly exceeded 23°C, with MHW peaks reaching 25°C in recent years (Fig. S1B), approaching the estimated T_lim_ for *P. scouleri*. Mass mortality events have already been reported for seagrass populations with comparable TSMs across both temperate and tropical regions (Marbà & Duarte, 2010; Moore *et al*., 2014; Thomson *et al*., 2015; Carlson *et al*., 2018), suggesting that *P. scouleri* may likewise be vulnerable to future climate-driven warming events. This risk is further supported by biogeographic projections predicting range contractions and genetic erosion at the southern edge of *Phyllospadix* spp., including northern Baja California populations, under future warming scenarios (Tavares *et al*., 2024).

Further supporting this concern, global-scale projections indicate that climate change may result in widespread reductions of seagrass biomass of up to 9.25%, with significant losses expected in warm-temperate and tropical regions (Gouvêa *et al*., 2025). This observation suggests the possibility that the growth suppression observed in *P. scouleri* under extreme warming could foreshadow broader declines in seagrass meadows as marine heatwaves become more frequent and intense. Similar thermal thresholds have driven long-term productivity declines in *Posidonia oceanica* (Litsi-Mizan *et al*., 2023), exemplifying the potential consequences of carbon imbalance events in seagrass systems.

Such losses are not merely reductions in biomass, they represent a weakening of critical ecosystem functions. As foundational species, seagrasses play a central role in nutrient cycling, carbon sequestration, shoreline stabilization, and habitat provision for diverse marine communities (Unsworth *et al*., 2022). The disruption of nitrogen assimilation observed in *P. scouleri*, particularly the suppression of nitrate reductase activity (NRA) during heatwaves, indicates that warming can compromise essential metabolic pathways that sustain productivity and their role as effective biofilters in nutrient-enriched coastal waters (Bonet-Melià *et al*., 2023). If MHWs continue to exceed physiological thresholds, the long-term resilience of these ecosystems could be undermined. This evidences not only the ecological vulnerability of surfgrass meadows, but also the urgent need to monitor key physiological indicators such as nitrogen metabolism, respiration and growth, which can signal early declines in the ecosystem services these habitats provide.

This study reveals *P. scouleri*’s vulnerability to temperatures representative of future MHWs. However, the long-term persistence of this species is also contingent on seasonal and life-stage variability in thermal tolerance (Beca-Carretero *et al*., 2021; Olsen *et al*., 2012; Guerrero-Meseguer *et al*., 2017), and potential acclimation through stress memory mechanisms (Nguyen *et al*., 2020; Pazzaglia *et al*., 2021, 2022; Stipcich *et al*., 2022b). Moreover, thermal tolerance may differ among populations inhabiting environments with contrasting temperature regimes, such as along depth gradients, where plants at shallower sites are exposed to warmer and more fluctuating conditions (Marín-Guirao *et al*., 2016). Conversely, additional stressors such as eutrophication, light reduction, and altered trophic interactions may compound thermal impacts and further constrain *P. scouleri*’s resilience (Hernán *et al*., 2017; Mvungi & Pillay, 2019; Vivanco-Bercovich *et al*., 2022; Pazzaglia *et al*., 2022). These complex interactions complicate predictions of future seagrass persistence and underscore the need for integrated, multi-stressor studies to better anticipate *P. scouleri*’s resilience under climate change.

Moving forward, research should examine thermal vulnerability across seagrass life stages, with particular attention given to germination and seedling development, which have been found to be more sensitive to environmental stress (e.g., Rinaldi *et al* 2023). In the context of surfgrasses, the broader impacts of MHWs on spatial distribution and long-term persistence remain uncertain, raising questions about whether warm-edge populations exhibit greater tolerance or heightened vulnerability (Bennet *et al*., 2022). Beyond plant-specific traits, future studies should also address how additional MHW characteristics, including duration, timing, and interval between events, shape physiological responses and recovery trajectories. Expanding such work across species and bioregions will facilitate the identification of generalizable thresholds of resilience and the differentiation of context-dependent responses.

## Author contributions

M.V.B., P.B.M., and J.M.S.G. planned and designed the research.

M.V.B., P.B.M., J.A.G.P., J.M.S.G., and J.M.G.C. conducted the fieldwork, experiments and collected the data.

M.V.B. and A.F.A. performed the laboratory analyses with input from P.B.M.

M.V.B. and J.M.S.G. analyzed data.

M.V.B. led the writing of the manuscript with substantial contributions from all co-authors. All authors discussed the results and approved the final version of the manuscript.

## Data availability

All primary data supporting this study are prepared for deposition in Dryad and will be publicly available upon article acceptance. The dataset will include raw and processed physiological measurements, experimental temperature records, and statistical scripts.

## Abbreviations

α: Photosynthetic efficiency
DP: Daily productivity
E_c_: Compensation irradiance
E_k_: Saturation irradiance
ETR: Electron transport rate
F_0_: Minimum fluorescence
F_m_: Maximum fluorescence
F_v_/F_m_: Maximum quantum yield
Gross-P_max_: Gross maximum photosynthetic rate
MHW: Marine heatwave
Net-P_max_: Net maximum photosynthetic rate
NPQ: Non-photochemical quenching
NRA: Nitrate reductase activity
NSC: Non-structural carbohydrates
R: Respiration rate
RGR: Relative growth rate
Φ_PSII_: Effective quantum yield.

---

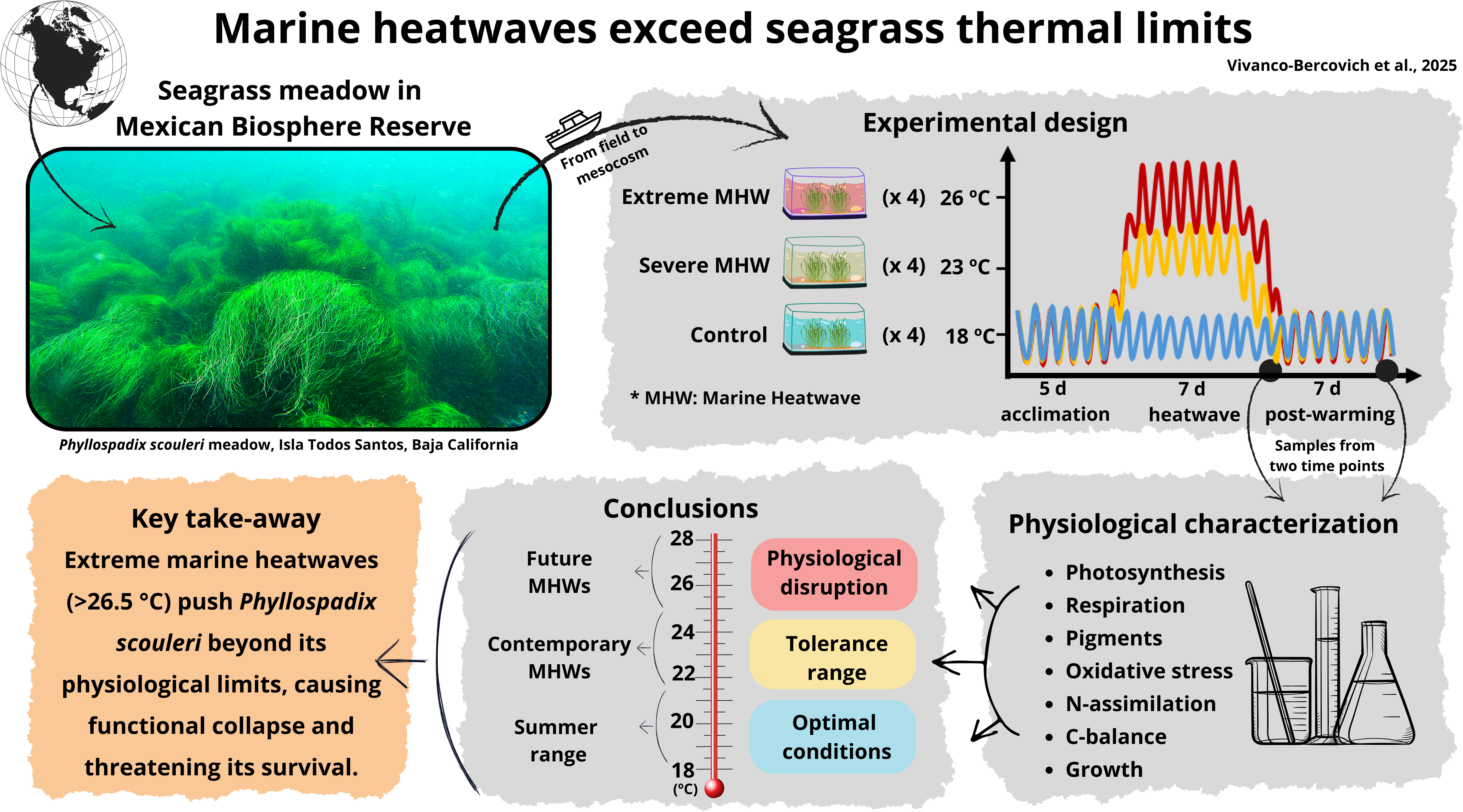

